# Acidotolerant soil nitrite oxidiser ‘*Candidatus* Nitrobacter laanbroekii’ NHB1 alleviates constraints on growth of acidophilic soil ammonia oxidisers

**DOI:** 10.1101/2024.07.06.601931

**Authors:** Linda Hink, Eleftheria Bachtsevani, Yiyu Meng, Christopher J. Sedlacek, Sungeun Lee, Holger Daims, Michael Wagner, Cécile Gubry-Rangin, Wietse de Boer, Christina Hazard, James I. Prosser, Graeme W. Nicol

**Affiliations:** CNRS, INSA Lyon, Université Claude Bernard Lyon 1, Ecole Centrale de Lyon, Ampère, UMR5005, 69134 Ecully, France; Institute of Microbiology, Leibniz University Hannover, Hannover, Germany; School of Biological Sciences, Cruickshank Building, University of Aberdeen, St Machar Drive, AB24 3UU, Scotland, UK; Centre for Microbiology and Environmental Systems Science, University of Vienna, Austria; Department of Microbial Ecology, Netherlands Institute of Ecology (NIOO-KNAW), PO Box 50, 6700AB Wageningen, the Netherlands; Soil Biology Group, Wageningen University and Research, PO Box 47, 6700AA Wageningen, the Netherlands

**Author notes:** Corresponding authors: Graeme W. Nicol and Linda Hink. Equal contribution (Linda Hink, Eleftheria Bachtsevani).

**Keywords:** *Nitrobacter*, nitrite-oxidising bacteria, ammonia-oxidising archaea, acidophilic, acidic soil, ammonia oxidation, nitrite oxidation, nitrification, cyanase

## Abstract

*Nitrobacter* strain NHB1 is a nitrite-oxidising bacterium previously co-enriched with the neutrophilic ammonia-oxidising bacterium *Nitrosospira* AHB1, a consortium that nitrifies under acidic conditions. Here we characterise the growth of isolated *Nitrobacter* strain NHB1 as a function of pH and nitrite (NO_2_^-^) concentration, and its influence on the activity of acidophilic soil ammonia-oxidising archaea (AOA) in co-culture. NHB1 is acidotolerant and grows optimally at pH 6.0 (range 5.0 – 7.5) at initial NO_2_^-^ concentrations of 500 µM. However, the optimum decreases to pH 5.0 at lower initial NO_2_^-^ concentrations closer to those found in soil, with detectable growth down to pH 3.5. NHB1 has a comparatively high affinity for NO_2_^-^ with an apparent-half-saturation constant (54 µM) one order of magnitude lower than its closest relative, the neutrophilic strain *Nitrobacter hamburgensis* X14. In co-culture, NHB1 enhances the growth of acidophilic AOA. Specifically, *Nitrosotalea devaniterrae* Nd1 and *Nitrosotalea sinensis* Nd2 are sensitive to NO_2_^-^-derived compounds and only oxidise ∼200-300 µM ammonia (NH_3_) in batch cultures. However, in co-culture with NHB1, pH ranges were lowered by ∼0.5 pH units and both strains could oxidise up to 2.7-2.9 mM NH_3_, only limited by buffering capacity. NHB1 possesses a cyanase facilitating reciprocal cross-feeding via generating cyanate-derived NH_3_ and utilising AOA-derived NO_2_^-^. Removal of NO_2_^-^ is likely crucial for nitrifier growth in acidic soils and this study highlights the importance of considering substrate and metabolic product concentrations when characterising physiology. Genome analysis reveals that NHB1 is distinct from validated species and the name ‘*Nitrobacter laanbroekii*’ is proposed.

## Main text

Microbially-mediated nitrification in soil is typically dominated by chemolithoautotrophs. Canonical ammonia-oxidising archaea (AOA) and bacteria (AOB) can perform the first step whereby ammonia (NH_3_) is oxidised to nitrite (NO_2_^-^) and is coupled to the activity of canonical nitrite-oxidising bacteria (NOB) which subsequently oxidise NO_2_^-^ to nitrate (NO_3_^-^). In acidic soils, ammonia oxidation activity was considered paradoxical due to the reduced availability of NH_3_ (*pK_a_* for NH_3_:NH_4_^+^ = 9.25). In addition, ammonia oxidisers (AO) are sensitive to NO_2_^-^-derived nitrous acid (HNO_2_) and associated decomposition products formed at low pH [1], necessitating the use of buffers or neutralising media to sustain growth. The subsequent discovery and cultivation of acidophilic and acidotolerant AO explained this paradox to a large extent [2–5], but mechanisms including ureolytic activity [6, 7] or growth within biofilms [8] or aggregates [9] also facilitate growth of neutrophilic nitrifiers under acidic conditions. As canonical AO are present in environments together with NOB, enabling mutualistic interactions [9, 10], the cooperative activity of NOB in soil is likely crucial for AO activity.

*Nitrobacter* populations are a major component of soil NOB communities. While most isolates are grown at neutral pH [11], acidophilic/acidotolerant strains have been reported [9, 12]. These include *Nitrobacter* NHB1 which was originally cultivated from pH 3.8 fertilised heathland soil together with the AOB *Nitrosospira* AHB1 [13]. Although neutrophilic, AHB1 was active down to pH 4 when grown with NHB1 [14] and neutrophilic *Nitrosospira*-like bacteria enriched from acidic forest soil grew at pH 4 when surrounded by *Nitrobacter*-like bacteria in aggregates [9]. Acidotolerant NOB may therefore have a role in protecting ammonia oxidisers in acidic soils by removing NO_2_^-^ before abiotic conversion to toxic compounds.

In this study we investigated the cell structure, genome content and growth characteristics of the isolate NHB1, including the effect of pH on its growth and a potential influence of NO_2_^-^ concentration in defining its pH range. As NO_2_^-^ does not typically accumulate in soil and isolated acidophilic AOA are particularly sensitive to NO_2_^-^-derived compounds in culture, we tested the hypothesis that NHB1 will positively impact the growth characteristics of *Nitrosotalea* strains in co-culture, which may reflect more accurately their ecophysiology *in situ*.

### Isolation and characterisation of NHB1

NHB1 was isolated from a cryopreserved co-culture of NHB1 and AHB1 by substituting NH_4_Cl in the growth medium with NaNO_2_ (Supplementary Methods). NHB1 is rod-shaped with typical stacked intracytoplasmic membranes (Fig. 1a) and possesses a 3.3 Mb chromosome and two plasmids (0.28 Mb and 0.21 Mb). The closest validated relative is the neutrophile *Nitrobacter hamburgensis* X14 [15] (Fig. 1b), sharing 99, 97, and 98% identity with 16S rRNA, nitrite oxidoreductase sub-unit alpha (NxrA) and beta (NxrB) encoding genes, respectively. High percentage identity of marker genes between *Nitrobacter* species is typically observed [16] and contrasts with NHB1 having a 24% smaller genome than *N. hamburgensis* X14 and sharing an average nucleotide identity of 92.4%. We therefore propose the following candidate species: *Nitrobacter laanbroekii* sp. nov. (laan.broek’i.i. N.L. gen. n. *laanbroekii*), named in honour of the Dutch microbiologist Hendrikus J. Laanbroek who was involved in its original cultivation and has made valuable contributions to understanding the ecology of NOB. In addition to expected core genes for carbon and energy metabolism, NHB1 possesses genes for dissimilatory sulphur oxidation, assimilatory nitrite reductase and utilisation of carbon monoxide as found for other *Nitrobacter* species [16].

**Figure 1.**
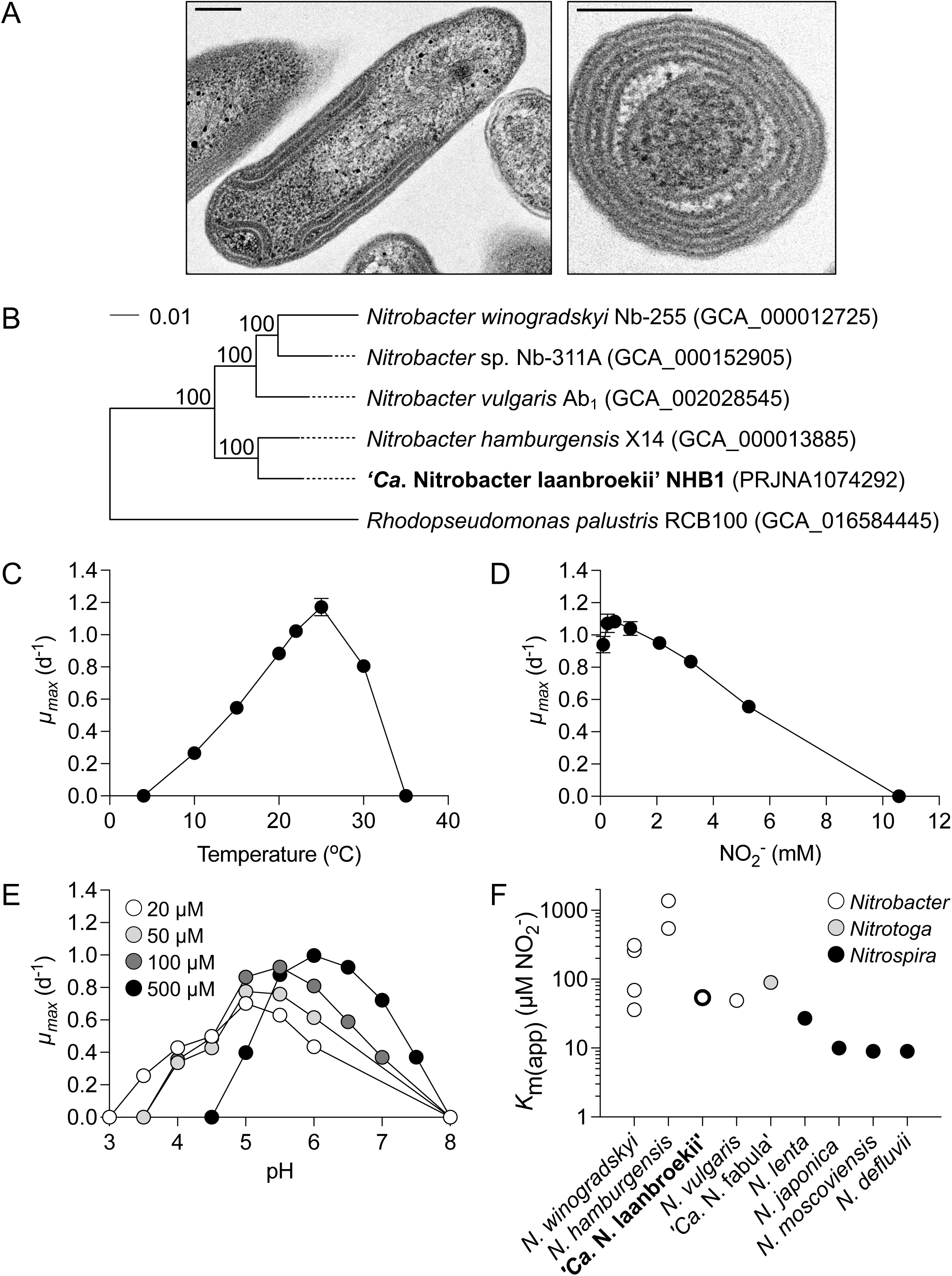
Characterisation of the isolated strain ‘*Ca.* Nitrobacter laanbroekii’ NHB1. (A) Transmission electron micrograph of cells in lateral and longitudinal orientation. Scale bars represent 0.2 µm. (B) Maximum likelihood phylogenomic tree of NHB1, selected *Nitrobacter* isolates and outgroup reference *Rhodopseudomonas palustris* (also of the *Nitrobacteraceae*) using 11,751 unambiguously aligned amino acid positions inferred from 63 single copy genes. The scale bar represents an estimated 0.01 changes per position and values at nodes describe percentage bootstrap support (100 replicates). The influence of (C) temperature, (D) initial nitrite concentration, and (E) pH on maximum specific growth rates (*µ_max_*) were determined with mean values and standard errors (mostly smaller than symbol size) from triplicate cultures plotted. (F) Apparent half-saturation constant (*K*_m(app)_) for NO_2_^−^ of ‘*Ca.* Nitrobacter laanbroekii’ NHB1 (mean value of four replicates) and various *Nitrobacter, Nitrotoga* and *Nitrospira* strains (Supplementary Methods).

Growth occurred with initial NO_2_^-^ concentrations up to >5 mM (inhibited at ∼10 mM) and temperatures ranging 10-30°C, with a maximum specific growth rate (*µ_max_*) of 1.17 d^-1^ (s.e.=0.05) at 25°C and 0.5 mM (Fig. 1c and 1d). As indicated from its previous enrichment, NHB1 is acidotolerant but *µ_max_* at different pH varied with initial NO_2_^-^ concentration. At 500 µM, growth occurred at pH 5.0-7.5 with optimal growth at pH 6.0 (Fig. 1e), but reduction of initial NO_2_^-^ concentration to 100 µM decreased optimal growth pH to 5.5 (*µ_max_* 0.93 d^-1^ (s.e.=0.01)) with further reductions in pH growth optimum to 5.0 at 50 and 20 µM (*µ_max_* 0.77 d^-1^ (s.e.=0.03) and 0.70 d^-1^ (s.e.=0.01), respectively). At the lowest NO_2_^-^ concentration, the limit for growth was extended to pH 3.5 (*µ_max_* 0.26 d^-1^ (s.e.=0.01)). Whole cell nitrite oxidation kinetics at 25°C and pH 6 were determined using substrate concentration-dependent oxygen microrespirometry (Fig. S1). An apparent-half-saturation concentration (*K*_m(app)_) of 54 µM NO_2_^-^ (s.e.=7) was higher than several canonical nitrite-oxidizing *Nitrospira* but within the lower range of those observed for other *Nitrobacter* strains, and one order of magnitude lower than *N. hamburgensis* X14 (544 µM) [17] (Fig. 1f).

### Influence of NHB1 on acidophilic AOA physiology

*Nitrosotalea devaniterrae* Nd1 and *Nitrosotalea sinensis* Nd2 are two AOA strains with pH optima ∼5.0 [2]. Although obligately acidophilic with adaptations to low pH, both are sensitive to NO_2_^-^ with concentrations as low as 10 µM reducing growth rates [2]. In standard batch cultures with 500 µM NH_4_^+^ and pH 5.2, yields of NO_2_^-^ are typically 200-300 µm (Fig. 2a) and one to two orders of magnitude lower than neutrophilic soil AOA [18, 19]. However, in contrast to growth in isolation, all 500 µM NH_4_^+^ was oxidised when grown in co-culture with NHB1 (Fig. 2b). The amount of NH_4_^+^ oxidised could be extended to 1.2 and 2.7-2.9 mM for both strains when 2-(N-morpholino)ethanesulfonic acid (MES) buffer concentrations were increased to 20 and 50 mM, respectively (Fig. 2c, Fig. S2), with growth inhibited when pH eventually decreased to <3.8 (Fig. S2). In the presence of NHB1, the pH range of both strains was extended down by ∼0.5 pH units and *µ_max_* increasing significantly at pH ≤5.0 (*P*=<0.05) (Fig. 2d). NHB1 possesses genes encoding enzymes that were previously demonstrated to facilitate mutualistic interactions with AO, including cyanase which produces NH_4_^+^ from cyanate and enables reciprocal cross-feeding [10]. While cyanate abiotically degraded rapidly to NH_4_^+^ in pH 5.2 medium (Fig. S3), NHB1 cyanase activity was the dominant mechanism generating ammonium at pH 6.0 in cultures supplemented with 0.05 or 0.5 mM cyanate (Fig. 2e). Although ≥0.5 mM cyanate can support other AOA or nitrifying co-cultures [10], this concentration inhibited both *Nitrosotalea* strains, with less than 5% of NOB-generated ammonium being oxidised through to NO_3_^-^ and with no accumulation of NO_2_^-^ (Fig. S4). However, Nd1 and Nd2 used the majority or all cyanate-derived NH_4_^+^ at 0.05 mM, respectively, a concentration higher than that typically found in soil where cyanase is continuously turned over [20].

**Figure 2.**
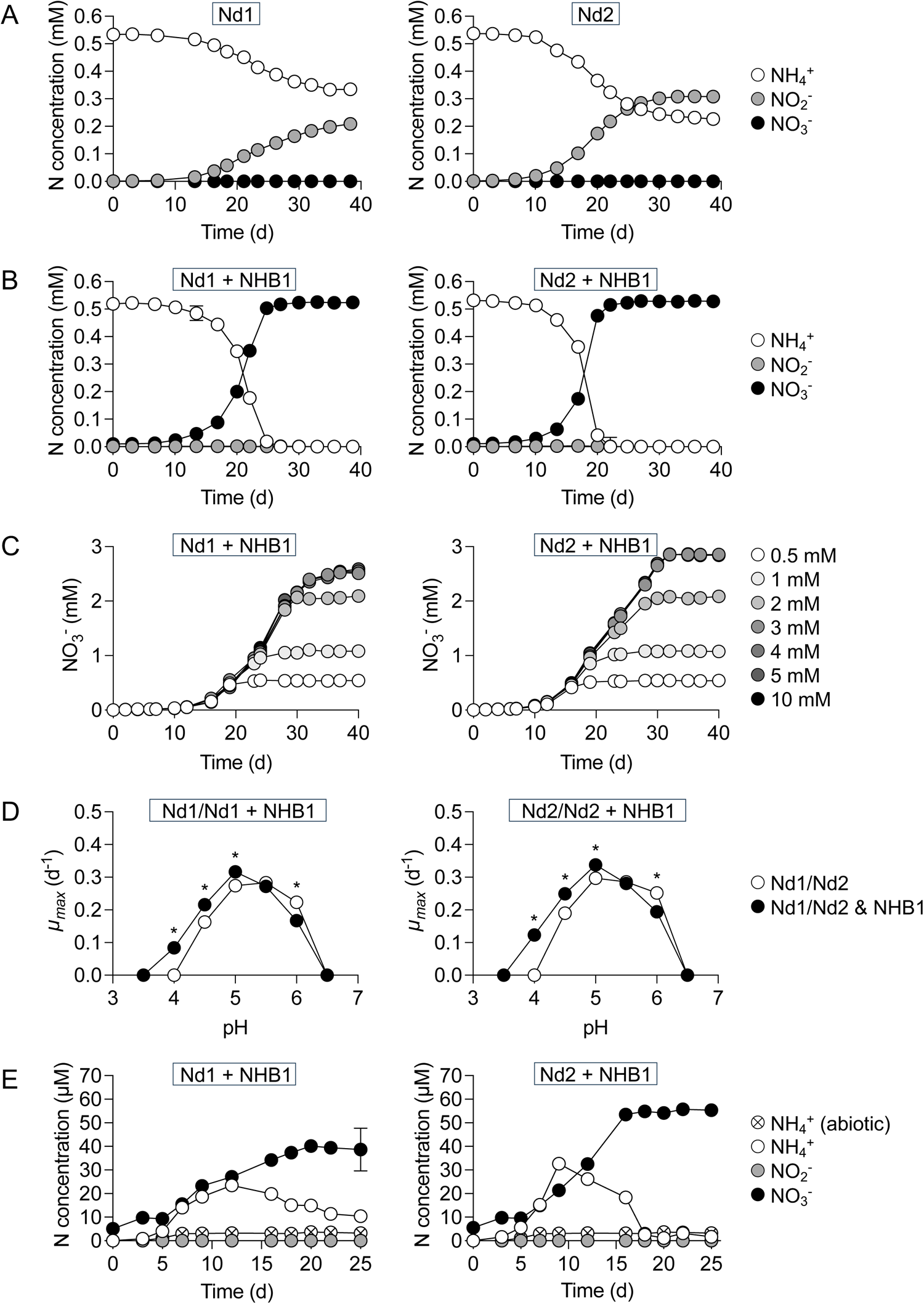
Influence of ‘*Ca.* Nitrobacter laanbroekii’ NHB1 on growth characteristics of acidophilic AOA *Nitrosotalea devaniterrae* Nd1 and *Nitrosotalea sinensis* Nd2 in batch culture. Mean values and standard errors’ (mostly smaller than symbol size) from triplicate cultures are plotted in each panel. (A) Growth of Nd1 and Nd2 grown in isolation and supplied with 0.5 mM NH_4_^+^. (B) Growth of Nd1 and Nd2 in co-culture with NHB1 supplied with 0.5 mM NH_4_^+^. (C) NO_3_^-^ production by co-cultures of Nd1 or Nd2 with NHB1 in medium with MES buffer concentration increased from 10 mM to 50 mM and 0.5, 1, 2, 3, 4, 5 or 10 mM NH_4_^+^. (D) The influence of NHB1 on the maximum specific growth rates (*µ_max_*) of Nd1 and Nd2 compared to growth in isolation at different pH. An asterisk highlights a significant difference in *µ_max_* (*p* = <0.05) for an individual pH (calculated using Student’s t-test). (E) Reciprocal cross-feeding at pH 6.0 between NHB1 and Nd1 or Nd2 with 50 µM cyanate supplied as an NH_4_^+^ source. NH_4_^+^, NO_2_^-^ and NO_3_^-^ concentrations were determined for all cultures. Decomposition of cyanate in the abiotic control produced <3.8 µM NH_4_^+^ during the period of incubation.

### Summary

‘*Ca*. Nitrobacter laanbroekii’ NHB1 is an acidotolerant NOB isolated from acidic soil with a relatively high affinity for NO_2_^-^. Growth in acidophilic consortia demonstrates that continued removal of NO_2_^-^ in acidic soil is likely crucial for sustained growth of AO and that consideration of substrate and metabolic product concentrations is essential when characterising physiology.

## Acknowledgements

This work was supported by an AXA Research Fund Chair awarded to GWN, the European Union’s Horizon WIDERA programme ‘ACTIONr’ under grant agreement No 101079299, and CGR was funded by a Royal Society University Research Fellowship (URF150571). The authors would like to thank Prof. Aharon Oren (Hebrew University of Jerusalem) for advice on etymology. We would also like to thank Elisabeth Errazuriz-Cerda and acknowledge the contribution of the CIQLE facility (a LyMIC member) at SFR Santé Lyon-Est (UAR3453 CNRS, US7 Inserm, UCBL) for transmission electron microscopy.

## Conflict of interest statement

The authors declare no conflicts of interest.

## Data availability

The genome sequence of ‘*Ca*. Nitrobacter laanbroekii’ NHB1 is available under NCBI BioProject accession number PRJNA1074292.

Supplemental Material

## Materials and methods

### Isolation and maintenance of ‘Ca. Nitrobacter laanbroekii’ NHB1 in culture

The co-culture of *Nitrosospira* sp. AHB1 and ‘*Ca*. N. laanbroekii’ NHB1 was obtained from cultures established in 1988 using pH 3.8 heathland soil (Hoorneboeg; 52°15’N, 5°10’E) under *Calluna vulgaris* (Heather) and *Deschampsia flexuosa* (Wavy hair-grass) vegetation [1, 2]. Cells from a cryostock were resuscitated by growing in 50 ml unbuffered medium with 2.5 mM NH_4_^+^ adjusted to pH 7. Specifically, the medium contained (L^-1^): KH_2_PO_4_ (0.1 g); NaCl (0.5 g); MgSO_4_·7H_2_O (0.04 g); CaCl_2_·2H_2_O (0.02 g); (NH_4_)SO_4_ (0.33 g); FeSO_4_·7H_2_O (2.46 mg); NaMoO_4_.2H_2_O (0.1 mg); MnCl_2_ (0.2 mg); Na_2_EDTA (3.31 mg); ZnSO_4_·7H_2_O (0.1 mg), CuSO_4_·5H_2_O (20 µg), and CoCl_2_ (2 µg) [1]. Cultures were incubated in the dark at 25°C without shaking. The same medium was used to purify ‘*Ca*. N. laanbroekii’ NHB1, except the pH was reduced to 5.5 and (NH_4_)_2_SO_4_ replaced with 500 µM NaNO_2_ (0.035 g L^-1^). Isolation was achieved by routine transfer (approximately weekly) of 1 ml exponentially growing culture into 50 ml fresh medium. The purity of ‘*Ca*. N. laanbroekii’ NHB1 was initially confirmed by the absence of ammonia oxidation activity when inoculated into fresh (NH_4_)_2_SO_4_-containing medium, loss of the band corresponding to the 16S rRNA gene amplicon of *Nitrosospira* sp. AHB1 in denaturing gradient gel electrophoresis (DGGE) analysis, and phase contrast microscopy (data not shown). Purity was confirmed by the absence of any contaminating DNA during genome sequencing and assembly.

The suitability of an acidophilic ammonia-oxidising archaea (AOA) medium for growing ‘*Ca*. N. laanbroekii’ NHB1 in potential co-culture experiments was assessed using NO_2_^-^ instead of NH_4_^+^ as an energy source. Specifically, acidic ‘freshwater medium’ (FWM) [3] consisted of NaCl (1 g L^-1^), MgCl_2_ (0.4 g L^-1^), CaCl_2_ (0.1 g L^-1^), KH_2_PO_4_ (0.2 g L^-1^), KCl (0.5 g L^-1^), 1 ml modifed non-chelated trace element solution [4], 1 ml 7.5 mM NaFeEDTA, 2 mM NaHCO_3_, and 500 µM NaNO_2_ (replacing 500 µM NH_4_Cl), was buffered with 10 mM 2-(N-morpholino)ethanesulfonic acid (MES) and adjusted to pH 5.5 before sterilisation by filtration using a GL45 bottle-top 0.2 µm filter unit (Nalgene, Rochester, USA). Successful growth of ‘*Ca*. N. laanbroekii’ NHB1 in this medium resulted in its subsequent use for routine growth and maintenance.

### Ammonia, nitrite and nitrate concentration measurements

All inorganic N concentrations were measured colorimetrically in 96-well plates using 50 µl of growth medium or adequate dilutions as described previously [5]. Briefly, NH_4_^+^ concentration was estimated using the indophenol method [6]. NO_2_^−^ and NO_3_^−^ concentrations were estimated using a method modified from Shinn [7] and Doane and Horwath [8]. NO_2_^-^ was detected by adding 60 μl of diazotising reagent (2.2 mM sulphanilamide in 3.3 M HCl) followed by 20 μl coupling reagent (0.12 mM N-(1-naphthyl)-ethylenediamine in 0.12 M HCl). NO_2_^−^ concentration was estimated immediately before reducing NO_3_^−^ to NO_2_^−^ by adding 20 μl vanadium chloride solution (4.5 mM vanadium(III) chloride in 1 M HCl), incubating for 90 min at 35°C in the dark and measuring NO_2_^−^ + NO_3_^−^ concentration.

### Characterisation of Ca. Nitrobacter laanbroekii NHB1

To determine temperature, pH and NO_2_^-^ concentration range, triplicate 50 ml cultures were statically incubated in sterile 100 ml culture bottles in the dark between 4 and 35°C, in medium adjusted to pH 3.0 to 8.0, and containing initial NO_2_^-^ concentrations between 20 µM and 10 mM, respectively. As NO_2_^-^ is known to degrade abiotically, particularly at pH lower than 5.5, triplicate sterile controls were also incubated with each treatment. All cultures were inoculated with 1% (vol/vol) early stationary phase culture that had consumed 500 µM NO_2_^-^. The growth of cultures was monitored via regular assessment of NO_2_^-^ concentrations. As NO_2_^-^ was stoichiometrically converted to NO_3_^-^ by ‘*Ca.* N. laanbroekii’ NHB1 (Fig. S5), measurement of NO_3_^-^ concentrations was not performed on a regular basis, but NO_3_^-^ production was calculated via NO_2_^-^ consumption after calculating the predicted abiotic degradation rate of NO_2_^-^, particularly under acidic pH (Fig. S6), enabling calculation of µ_max_ during the exponential growth phase of the cultures (Fig. S7).

### Growth of Nitrosotalea strains in isolation or co-culture with ‘Ca. Nitrobacter laanbroekii’ NHB1

Cultures of isolated *Nitrosotalea devaniterrae* Nd1 and *Nitrosotalea sinensis* Nd2 or stable co-cultures of *N. devaniterrae* Nd1 or *N. sinensis* Nd2 with ‘*Ca*. N. laanbroekii’ NHB1 were established by inoculating 1 ml of early stationary phase cultures into 50 ml standard FWM medium for acidophilic AOA (pH 5.2) containing 500 µM NH_4_Cl in sterile 100 ml culture bottles. While *N. devaniterrae* Nd1 and ‘*Ca*. N. laanbroekii’ NHB1 have similar temperature ranges and growth optima, *N. sinensis* Nd2 grows optimally ∼35°C [9] and above the temperature range of ‘*Ca*. N. laanbroekii’ NHB1. A temperature of 25°C was therefore used for isolation vs co-culture experiments. Growth of (co-)cultures was monitored via measurement of NH_4_^+^, NO_2_^-^ and NO_3_^-^ concentrations until stationary phase. The effect of pH and substrate concentration was determined in 30 ml sterile plastic Universal bottles (Greiner Bio-One, Les Ulis, France) containing 20 ml of medium. Co-cultures can be maintained indefinitely by transfer (2% vol/vol) every 2-3 weeks into sterile medium.

In co-culture experiments to examine the potential cyanase activity of NHB1, ammonium was substituted with 50 or 500 µM cyanate. In experiments examining yields of NO_3_^-^ with higher NH_4_^+^ concentrations, despite the absence of NO_2_^—^derived acids in co-culture, pH decreased with growth and higher concentrations of MES (both 20mM and 50 mM) were required to sustain growth after consumption of ∼800 µM NH_4_^+^.

### Determination of substrate kinetics

Cellular nitrite oxidation kinetics were determined from instantaneous substrate-dependent oxygen uptake measurements as previously described using a multiple injection method [10, 11]. Briefly, measurements were performed in a water bath at 25°C with a microrespirometry (MR) system, equipped with a PA2000 picoammeter and a 500 μm tip diameter OX-MR oxygen microsensor (Unisense, Aarhus, Denmark), polarized continuously for at least 24 h before use. Active NHB1 cells were taken from early stationary phase cultures soon after substrate depletion or harvested and concentrated (6000 x *g*, 10 min, 20 °C) from NO_2_^-^ replete active cultures using Amicon Ultra-15 10 kDa cut-off centrifugal filter units (Merck Millipore, Darmstadt, Germany). Concentrated cells were washed with and resuspended in substrate-free medium (pH 6) prior to MR measurements. Whole cell activity rates of NHB1 were fit with the Michaelis-Menten model to derive the apparent whole cell reaction half saturation concentration (*K*m_(app)_; µM NO_2_^-^). The *K*m_(app)_ of *‘Ca.* N. laanbroekii’ NHB1 was compared with other nitrite oxidiser strains using previously published values [12–15]

### Transmission electron microscopy

NHB1 cells were recovered from 1 L of late exponential culture by filtering onto a 0.2 µm mixed cellulose ester filter before resuspending cells in 2 ml fresh medium and pelleting by centrifugation at 14,000 x *g* for 40 min. One volume (500 µl) 4% glutaraldehyde (v/v) was added to ∼500 µl medium overlying the pellet and stored at 4°C overnight. Cells were then post-fixed with 1% OsO_4_ in 0.3 M cacodylate buffer (pH 7.4) for 1 h at 4°C before ethanol dehydration and transfer to propylene oxide. Cells were embedded in Epon epoxy resin and inclusion obtained by polymerisation at 60°C for 72 h. Ultra-thin sections (100 nm) were cut using a UC7 ultramicrotome (Leica, Nanterre, France), mounted on 200 mesh copper grids (EMS, Hatfield, USA) and contrasted with uranyl acetate and lead citrate. Sections were examined with a JEM-1400 120 kV transmission electron microscope (Jeol, Tokyo, Japan) equipped with a Orius 1000 camera (Gatan, Pleasanton, USA) in the wide-field position and Digital Micrograph software (Gatan) at the Centre d’Imagerie Quantitative Lyon-Est, Université Claude Bernard Lyon 1.

### Genome sequencing, assembly and annotation

DNA was extracted from pelleted cells using a standard SDS buffer and phenol: chloroform:isoamyl alcohol chemical lysis method [16] and sequenced using in-house MiSeq (Illumina, San Diego, USA) and MinION (Oxford Nanopore, Oxford, UK) platforms. A library for MiSeq paired-end sequencing was prepared using a Nextera XT kit, according to the manufacturer’s instructions (Illumina) and sequencing performed with a V2 kit, producing 20.1 million reads with an average length of 243.4 bp after quality trimming performed using TrimGalore (https://github.com/FelixKrueger/TrimGalore). For MinION sequencing, DNA was sheared to approximately 8 kb using a g-TUBE (Covaris, Brighton, UK) and a library prepared with the SQK-MAP006 kit, according to the manufacturer’s instructions (Oxford Nanopore). Sequencing on a MinION flow cell produced 39,064 reads with an average length of 6,739 bp. The genome was assembled using Unicycler [17] with default settings for both MinION and Illumina data. Gene prediction was performed using Prodigal version 2.6.3 [18] and annotation performed using Diamond BLASTp v0.8.36 (e-value <10^−5^) [19] with the NCBI nr database release 244 [20]. Genome quality was assessed using CheckM [21], estimating 99.1% completeness and 0.34% contamination, and GToTree [22], detecting 100% of expected single copy genes with 0% redundancy. Average nucleotide identity (ANI) between different *Nitrobacter* strains was assessed using reciprocal best hits (two-way ANI) (http://enve-omics.ce.gatech.edu/ani/) [23].

### Phylogenomic analysis

Single copy genes present in all compared genome sequences were identified and aligned using GToTree [22] before manual refinement. Maximum likelihood analysis was performed on unambiguously aligned concatenated protein sequences (11,751 amino acid positions inferred from 63 single copy genes) using PhyML [24] with automatic model selection (Q.plant with FreeRate variation (four rates) across sites) and bootstrap support (100 replicates).

---

**Figure S1.**
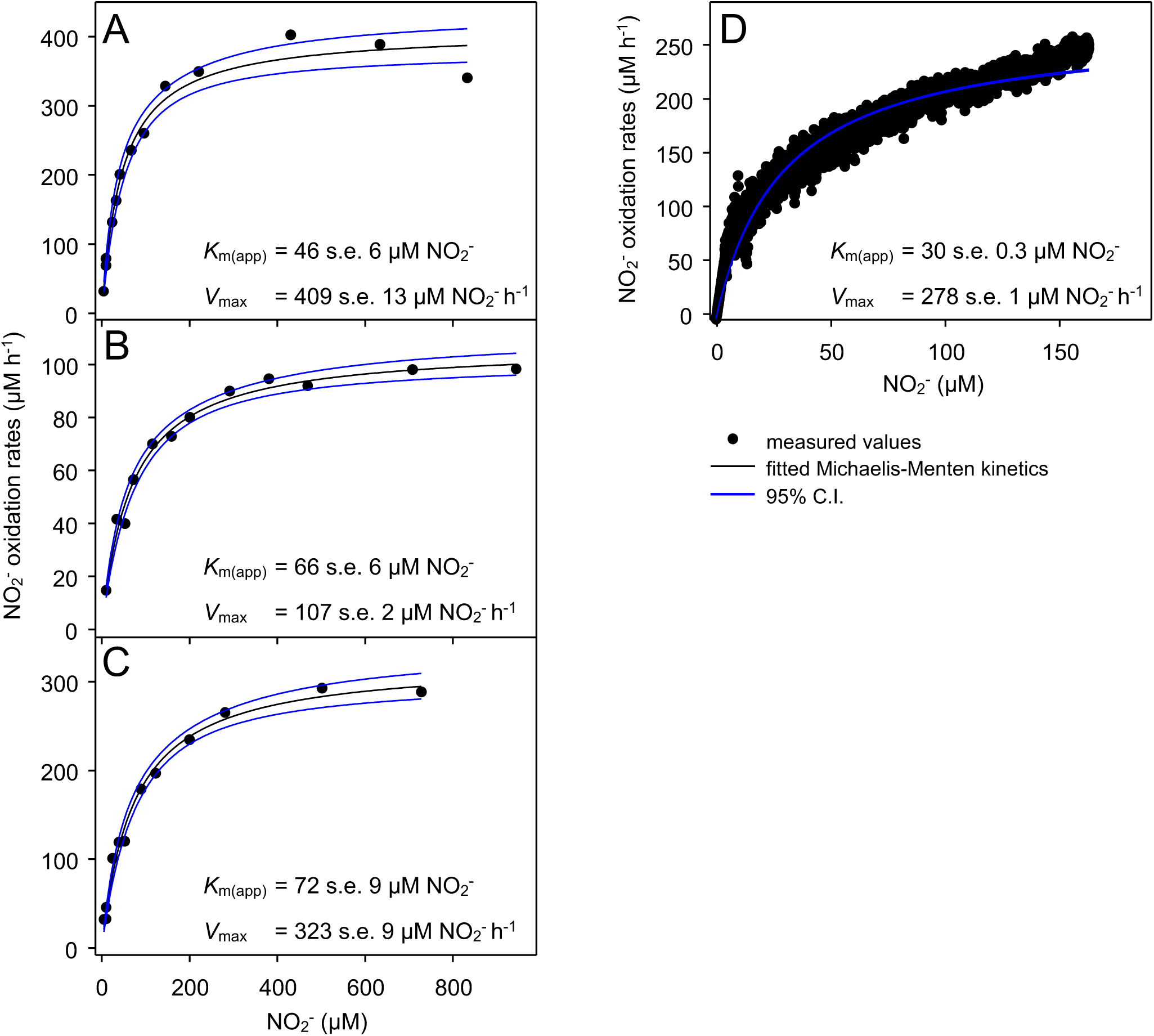
Apparent half-saturation (*K*_m(app)_) and maximum oxidation rates (*V*_max_) for NO_2_^-^ calculated using oxygen microrespirometry (MR) after fitting data of oxidation rate vs NO_2_^-^ concentration to the Michaelis–Menten model with 95% confidence intervals (C.I.) plotted. Three experimental replicates measured oxidation rates after injection varying concentrations of NO_2_^-^ (A,B,C) into the MR chamber and one replicate (D) used a single injection of 250 µM. Biomass was not measured and *V*_max_ was therefore not normalised to cell numbers or protein content and varies substantially amongst replicates.

**Figure S2.**
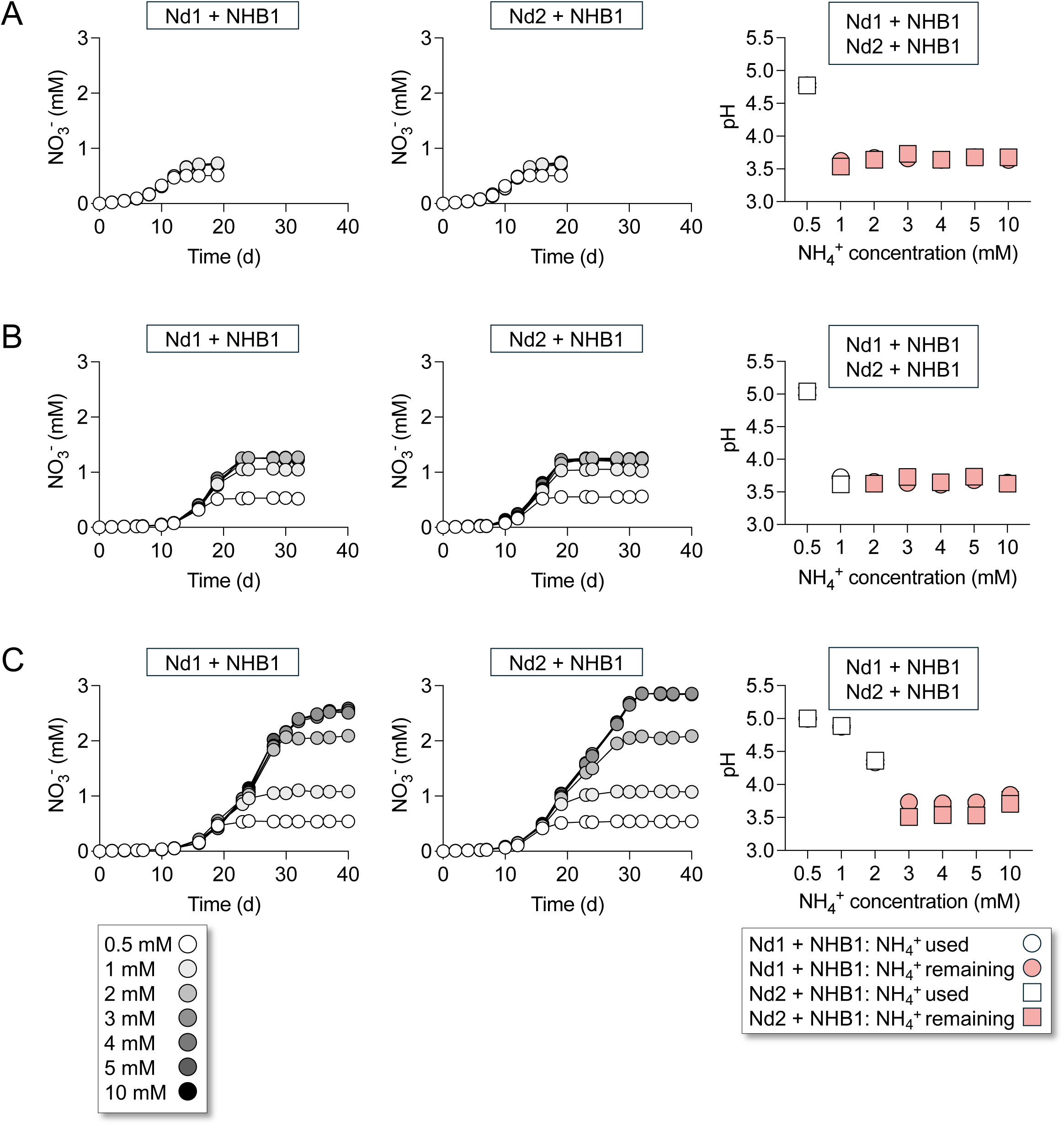
Effect of MES buffer concentration (**A** 10 mM, **B** 20 mM, **C** 50mM) on the growth of co-cultures of ‘*Ca.* Nitrobacter laanbroekii’ NHB1 and *Nitrosotalea devaniterrae* Nd1 or *Nitrosotalea sinensis* Nd2 in medium with an initial pH 5.2 and different NH_4_^+^ concentrations (0.5, 1, 2, 3, 4, 5 and 10 mM). NO_3_^-^ concentrations were determined through to stationary phase and pH measured at the last time point. The presence of remaining NH_4_^+^ at the end of the stationary phase is described and demonstrates inhibition by decreasing pH rather than substrate limitation. Mean values and standard errors (mostly smaller than symbol size) from triplicate cultures are plotted in all panels.

**Figure S3.**
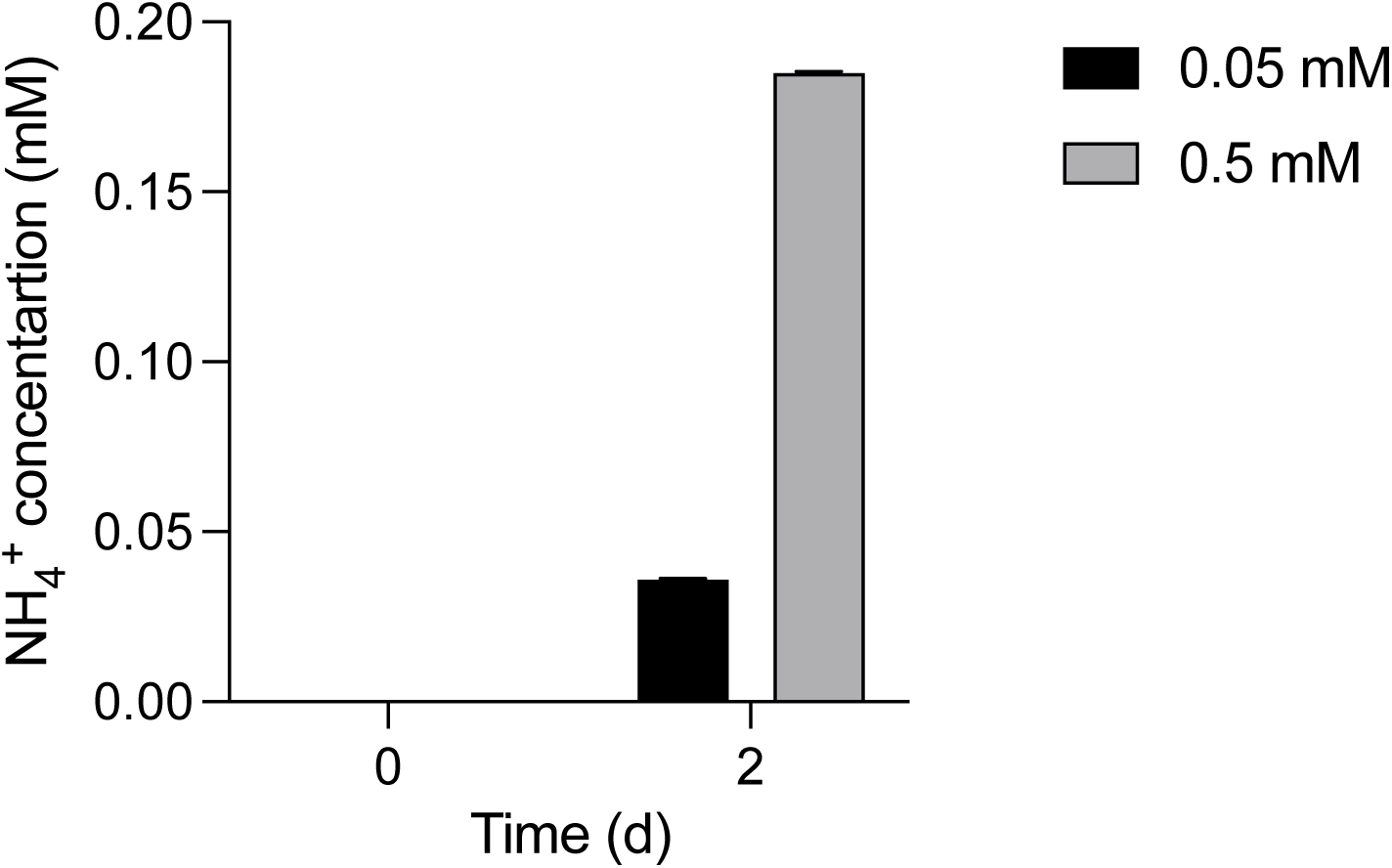
Production of NH_4_^+^ from abiotic decomposition of cyanate in pH 5.2 ‘freshwater medium’ amended with 0.05 mM or 0.5 mM after two days incubation at 25°C.

**Figure S4.**
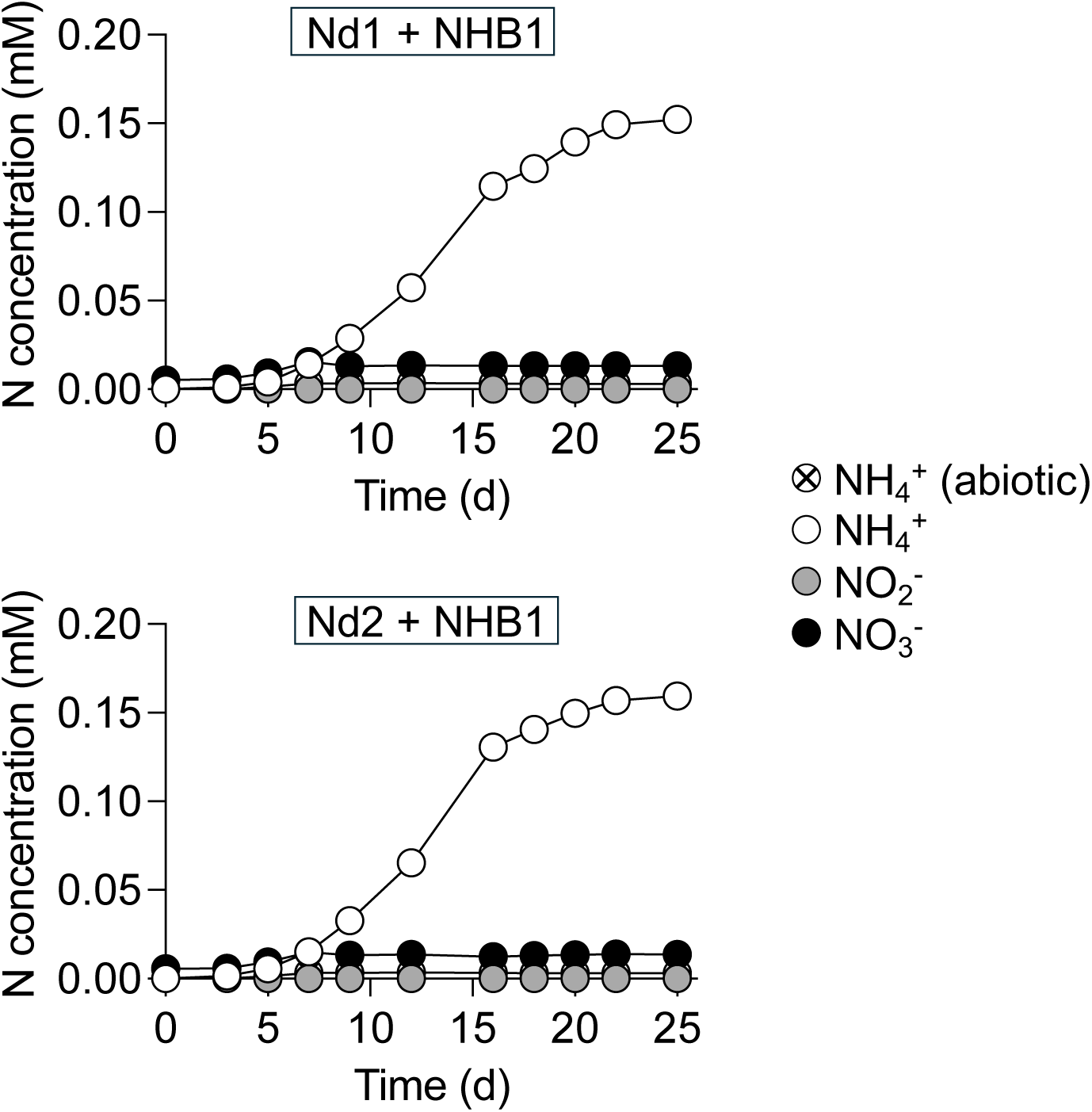
Inhibition of acidophilic AOA in ‘freshwater medium’ batch cultures (pH 6.0) amended with 0.5 mM cyanate. Cultures were inoculated with a previously established co-culture of ‘*Ca.* Nitrobacter laanbroekii’ NHB1 and *Nitrosotalea devaniterrae* Nd1 or *Nitrosotalea sinensis* Nd2. Less than 5% of NOB-generated ammonium was oxidised through to NO_3_^-^ with no accumulation of NO_2_^-^. Decomposition of cyanate in the abiotic control produced <3.5 µM NH_4_^+^ during the period of incubation. Mean values and standard errors (mostly smaller than symbol size) from triplicate cultures are plotted.

**Figure S5.**
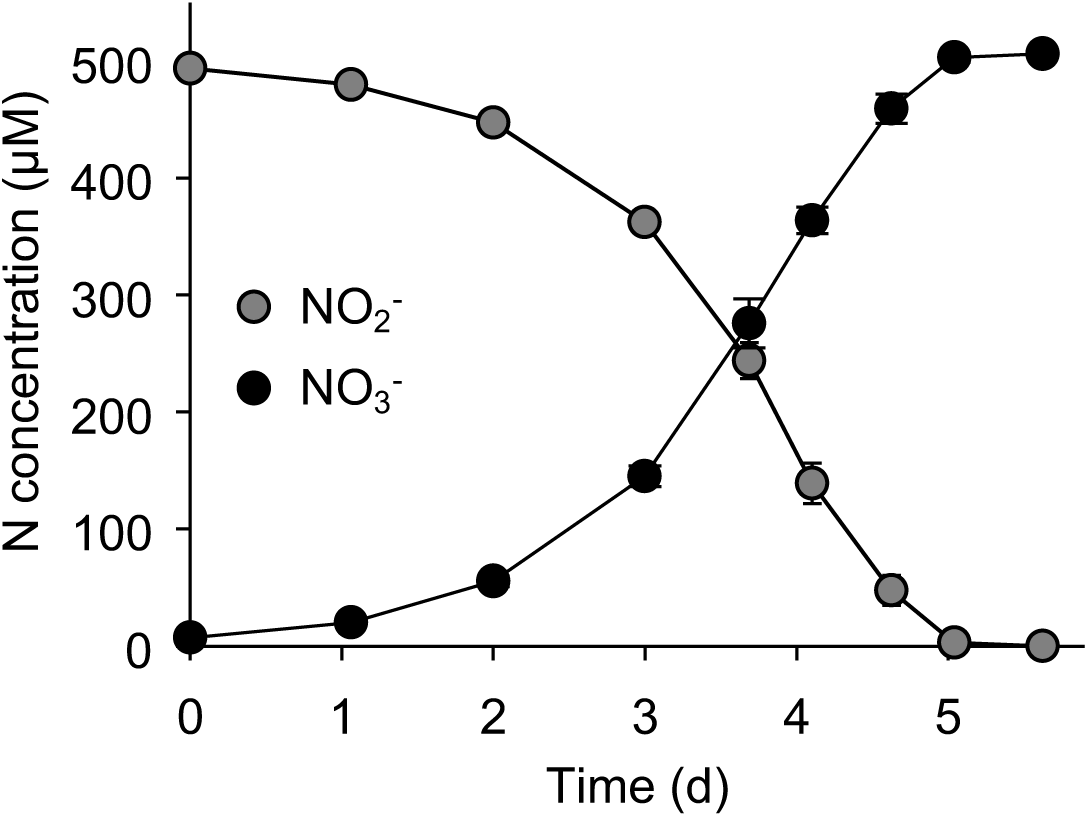
Stoichiometric relationship of NO_2_^-^ consumption and NO_3_^-^ production during growth of ‘*Ca.* Nitrobacter laanbroekii’ NHB1. The medium was adjusted to pH 5.5 with an initial NO_2_^-^ concentration of 500 µM and inoculated with 1% early stationary culture (vol/vol) and incubated at 25°C in the dark. Plotted values are the mean and standard errors of NO_2_^-^ and NO_3_^-^ concentrations of triplicate cultures. The confirmed stoichiometry between NO_2_^-^ consumption and NO_3_^-^ production enabled the use of NO_2_^-^ concentrations to infer NO_3_^-^ concentrations in subsequent culture characterisation experiments.

**Figure S6.**
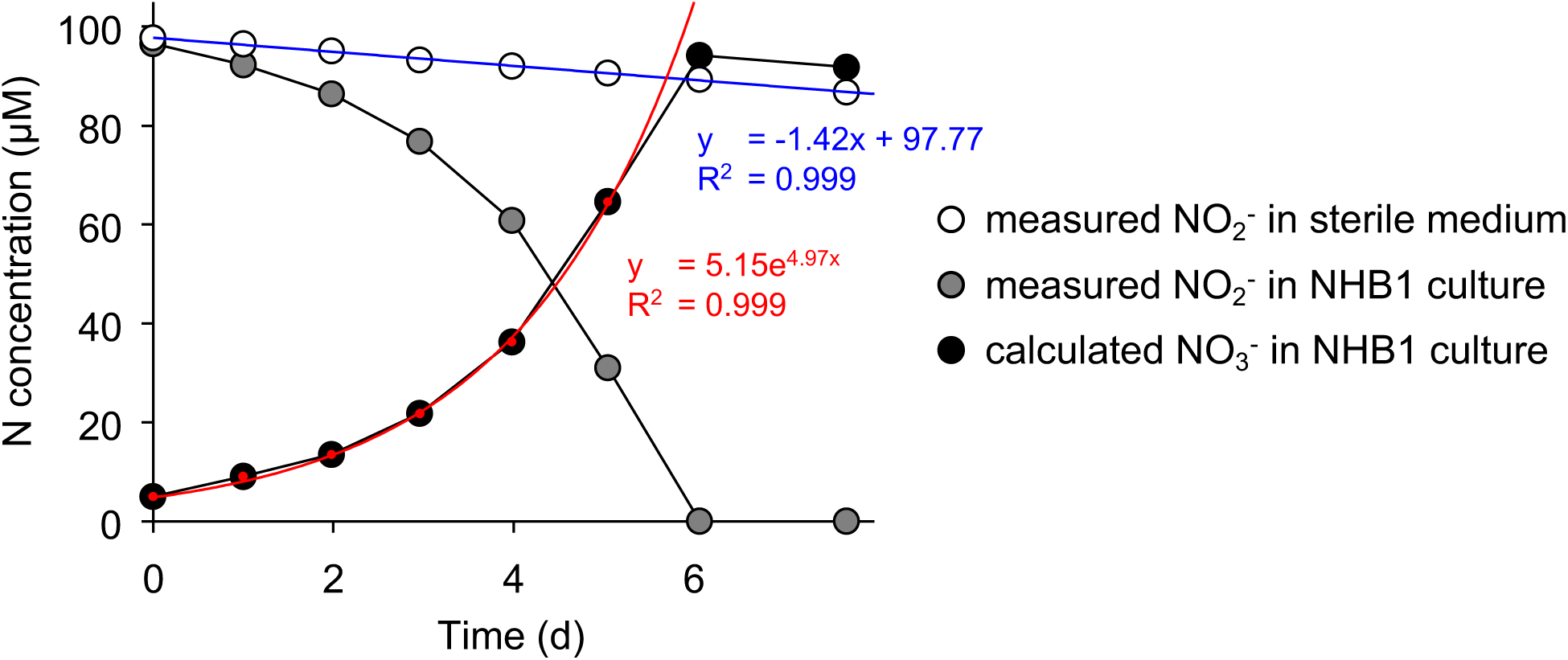
An example of assessing *µ_max_* of ‘*Ca.* Nitrobacter laanbroekii’ NHB1 when grown at pH 4.5 with an initial 100 µM NO_2_^-^ concentration. Medium was inoculated with a 1% transfer (vol/vol) of an early stationary culture that had consumed 500 µM NO_2_^-^. Sterile medium served as control to monitor abiotic NO_2_^-^ degradation following a linear decline over time (blue regression and equation). Measured NO_2_^-^ concentrations in the growing culture and the abiotic degradation rate NO_2_^-^ in the sterile control was used to calculate predicted NO_3_^-^ concentrations. An exponential curve was then fitted to the NO_3_^-^ data during exponential growth of the culture (red circles, regression and equation). The parameter in the exponent of the equation corresponds to the *µ_max_* value of the culture.

**Figure S7.**
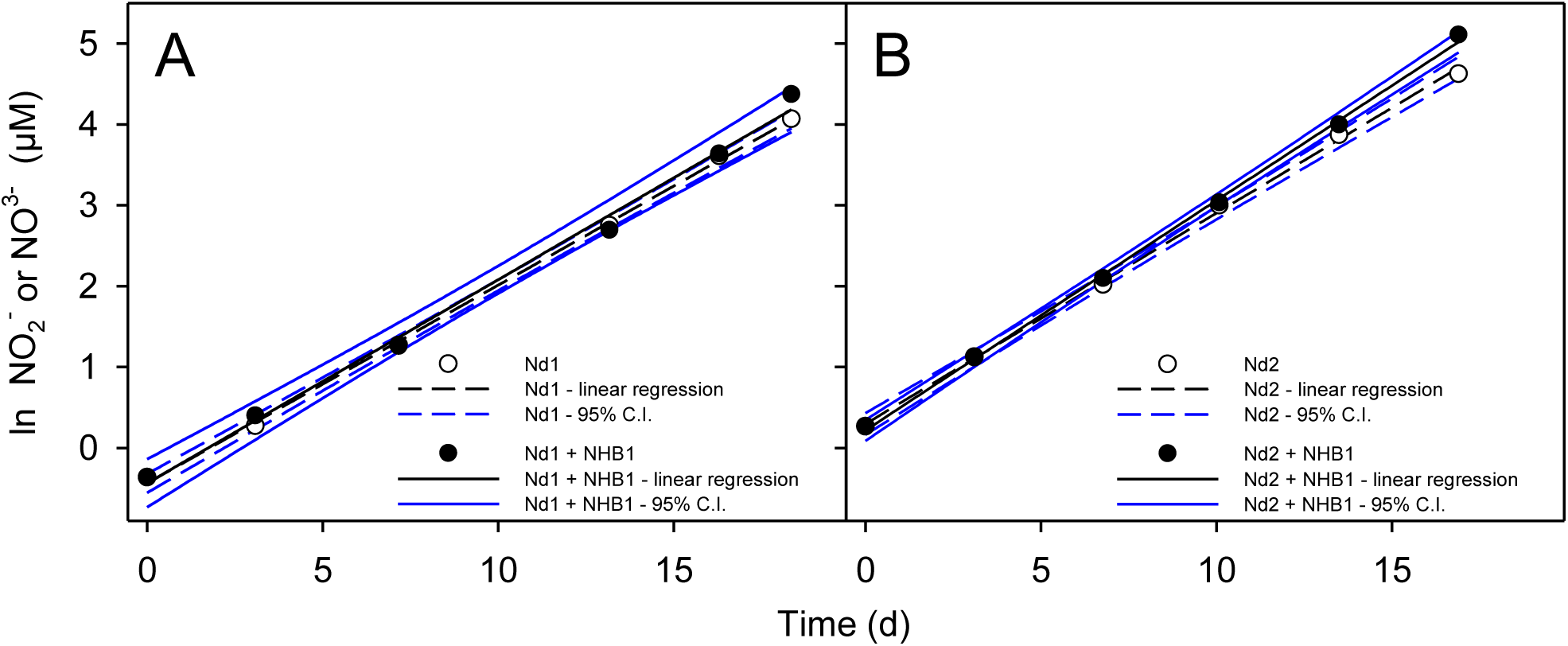
Examples of plots used for calculating maximum specific growth rates of *Nitrosotalea devaniterrae* Nd1 and *Nitrosotalea sinensis* Nd2 (assessed via NO_2_^-^ production) or in co-culture with ‘*Ca.* Nitrobacter laanbroekii’ NHB1 (assessed via NO_3_^-^ production) in medium supplied with 500 µM NH_4_^+^. Linear regression and 95% confidence intervals (C.I.) were calculated from natural logarithm (ln) values of NO_2_^-^ concentrations during the exponential phase of growth.

## Notes

### Competing Interest Statement

The authors have declared no competing interest.

